# Recurrent breakpoints in the *BRD4* locus reduce toxicity associated with gene amplification

**DOI:** 10.1101/2024.07.31.606009

**Authors:** Jeremiah Wala, Simona Dalin, Sophie Webster, Ofer Shapira, John Busanovich, Rameen Beroukhim, Pratiti Bandopadhayay, Veronica Rendo

## Abstract

Structural variants (SVs) represent a mechanism by which cancer cells activate oncogenes or disrupt the function of genes with tumor suppressor roles. A recent study by the PCAWG Consortium investigated structural variants in 30 tumor types, identifying focal rearrangements in the oncogenic BRD4 gene in ovarian, endometrial and breast cancers. These rearrangements resulted in decreased BRD4 expression despite increased copy number, suggesting a novel mechanism to finetune gene over-expression. In this study, we show that focal deletions of BRD4 disrupt genomic regulatory regions and impact gene isoform expression in breast and ovarian tumors when compared to their expression across normal tissues. To determine the functional impact of these concomitant amplification and focal deletion events, we first leveraged open-reading-frame (ORF) screen data from 16 cancer cell lines, where we observed that overexpression of BRD4-long and BRD4-short isoforms is toxic for cancer cell growth. We confirmed these results in OVSAHO ovarian cancer cells, where overexpression of both isoforms significantly reduced tumor growth. Next, we mimicked the focal deletions occurring in BRD4 regulatory regions by CRISPR-Cas9 technology, and observed that their depletion functionally ablates tumor cell growth. We finally show that these focal deletions rescue ovarian carcinoma cells from the toxicity effects associated with gene overexpression, suggesting that global BRD4 gene expression levels must be fine tuned to ensure proper cancer cell proliferation. Our study provides experimental evidence for BRD4 deletions constituting the first example of a driver SV alteration reducing toxicity in cancer, therefore expanding the landscape of cancer progression mechanisms.

## Introduction

Large-scale whole genome sequencing efforts have increasingly highlighted the role of genomic rearrangements in driving cancer progression. These structural variants (SVs) involve the ligation of genomically distant DNA regions, and can result in deletions, inversions, duplications, or more complex events. SVs can affect gene expression by altering copy number status, nearby regulatory elements, or by enabling novel fusion products^1–5^. Altogether, SVs represent a common mechanism by which cancer cells activate oncogenes (e.g. formation of *IGH-BCL2* and *KIAA1549-BRAF* fusions) or disrupt the function of tumor suppressor genes (e.g. *TP53*, *PTEN*).

A recent study from the Pan-Cancer Analysis of Whole Genomes (PCAWG) Consortium described the landscape of structural variants commonly occurring in 30 tumor types by analysis of 2,658 genomes^3^. From this effort, focal rearrangements within the oncogenic bromodomain-containing protein 4 (*BRD4*) in chromosome 19p were detected in a fraction of ovarian (n=8) and breast (n=7) cancers with amplification of this particular locus. The breakpoints identified at this locus resulted in small, focal deletions overlapping with intron 1 and often exon 1 of *BRD4*. Interestingly, amplified tumors exhibiting these rearrangements had constant or decreased levels of *BRD4* gene expression relative to their copy number. This was a striking observation, as increases in copy number for most genes in the genome tend to translate into increased gene expression^6^. This disruption in gene-dosage was hypothesized to constitute a novel mechanism to finetune gene over-expression in areas subject to *BRD4* genomic amplification. These *BRD4* deletions appear to be the first example of a driver SV alteration that reduces toxicity in cancer, but the functional impact of these focal deletions remains unknown.

BRD4 is a member of the bromodomain and extra-terminal domain family of proteins, able to bind acetylated histones to regulate gene transcription. Key biological roles include maintenance of chromatin structure to ensure epigenetic memory after mitosis, and transcriptional regulation of signal-inducible genes by association with the positive transcription elongation factor (p-TEFb) complex and RNA polymerase II^7–9^. *BRD4* is commonly altered in human cancers, and significant amplification is observed together with the known oncogene cyclin E1 (*CCNE1*) in both ovarian and breast carcinomas^10,11^.

In this study we sought to determine the impact of *BRD4* recurrent focal deletions on gene expression, with a particular focus on how isoform ratio is altered in *BRD4*-amplified ovarian and breast tumors. We additionally leverage functional genomic tools to assess the cellular effect of dysregulated *BRD4* expression across multiple human cancer cell lines. This validates the identified focal deletions in an ovarian cell line model as a mechanism to sustain cellular proliferation while dampening the toxicity effects associated with initial gene amplification. Our findings provide the first experimental evidence for a novel mechanism by which genetic alterations drive cancer.

## Results

### Significantly recurrent breakpoints result in focal deletion of *BRD4* regulatory regions

To better understand breakpoint distribution at the *BRD4* locus, we first accessed whole-genome sequencing data from the ICGC PCAWG cases where recurrent rearrangements were originally identified^3^. We looked at breakpoint densities present in all PCAWG cases, and confirmed that there is an enrichment of breakpoints within a small region of *BRD4*, typically ranging from exon 1 to intron 1 of the gene (**Figure 1A**). We were further able to identify similar patterns of rearrangements when comparing these samples to an additional breast cancer dataset^12^ (n=560 whole-genome sequences) (**Figure 1A**), supporting the notion that this may constitute a mutational event with functional implications in cancer.

**Figure 1.**
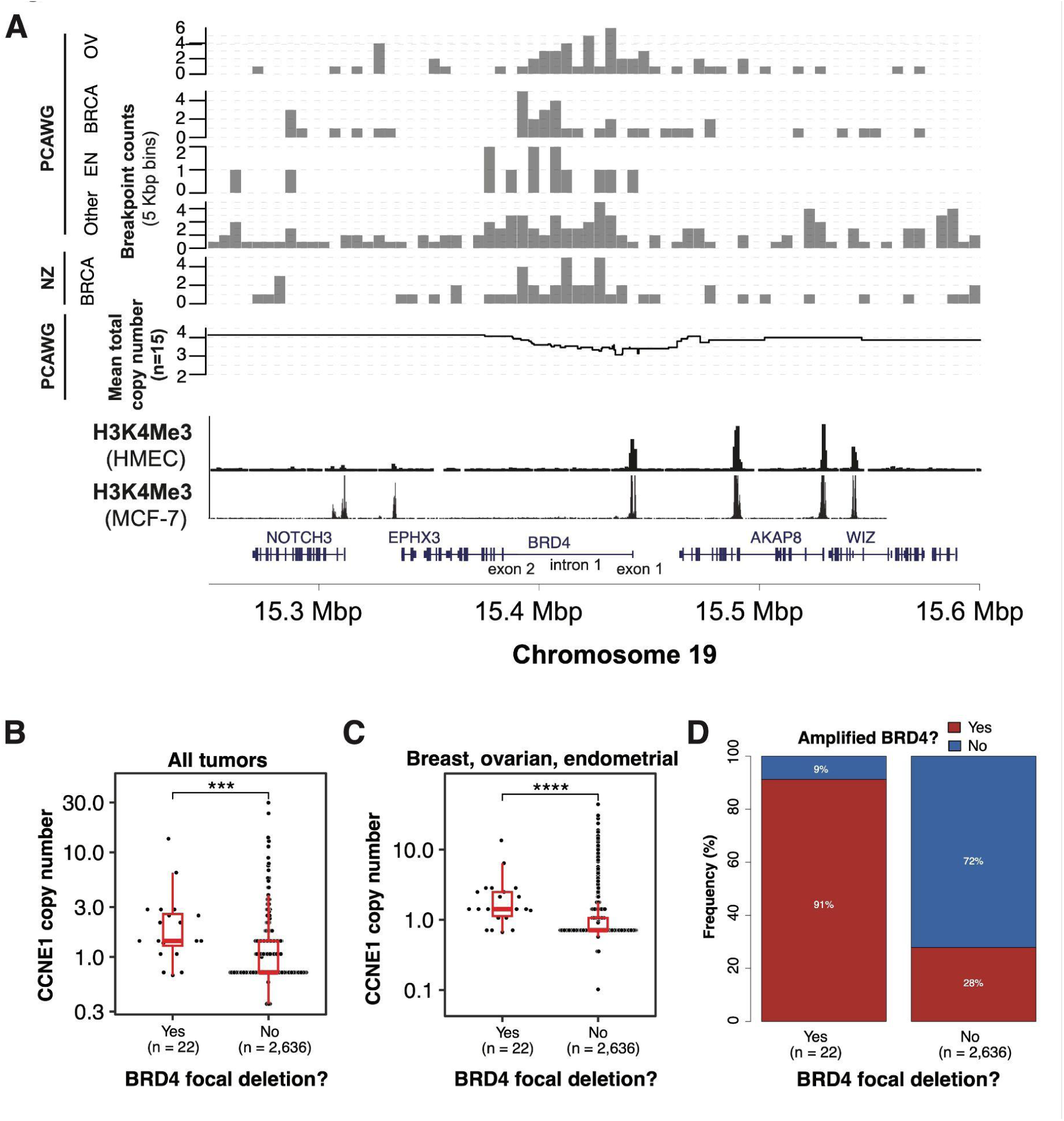
Landscape of focal deletions spanning the *BRD4* locus. **(A)** Top: Breakpoint densities for PCAWG breast (BRCA), endometrial (EN) and ovarian (OV) cancers exhibiting rearrangements at the *BRD4* locus, compared to an independent breast cancer dataset (Nik-Zainal et al.); Middle: Somatic copy-number alterations (SCNAs) for pooled breast, ovarian and endometrial cancer samples with *BRD4* focal deletions; Bottom: Alignment to H3K4me3 from breast (MCF-7) and ovarian (HMEC) cell lines. Genomic track: Gene exons and intron as defined in RefSeq^18^ for hg19. **(B)** Total somatic copy-number at *CCNE1* locus (chromosome 19) between tumors with or without *BRD4* focal deletions and **(C)** only breast, ovarian and endometrial tumors; Wilcoxon rank-sum test. **(D)** Proportion of tumors with or without amplifications spanning the *BRD4* locus, comparing those with or without a concomitant *BRD4* focal deletion.

To understand the effect of these recurrent breakpoints, we next analyzed the PCAWG somatic copy-number alterations (SCNA) data to look for SCNA events at the *BRD4* locus corresponding to the recurrent breakpoints. This revealed recurrent focal deletions (< 100 Kbp) in *BRD4* in 15 tumors, including 4 breast cancers, 6 ovarian, 3 endometrial and 2 colorectal cancers (**Extended Figure 1)**. We also identified an additional 12 cases involving a copy-loss of *BRD4* relative to the neighboring *NOTCH3* gene (1 breast, 3 ovarian, 2 endometrial, 3 pancreatic adenocarcinoma, 1 pancreatic endocrine, 1 lung squamous, and 1 hepatocellular cancers; **Extended Figure 1**). The total copy-number in genes adjacent to the *BRD4* locus tended to be higher in samples with focal *BRD4* deletions than in samples without *BRD4* deletions (*NOTCH3* median total copy-number: 4.7 vs. 2.4; p < 3 x 10^-11^, Wilcoxon-rank test; **Figure 1B**). Finally, we identified an additional 5 samples (3 breast and 2 ovarian) with breakpoints within the recurrent region of *BRD4*, but with no corresponding SCNA. On further inspection of the raw read-depth signal, these samples each exhibited a read-depth drop within the boundaries of the breakpoints, and with breakpoint orientations consistent with a focal deletion. Thus, we conclude that these also represent focal *BRD4* deletions given the commensurate read-depth and breakpoint signal, despite not having a sufficiently strong signal with read-depth alone to trigger an SCNA call (**Extended Figure 2**).

The enrichment for focal BRD4 deletions in breast, ovarian and endometrial cancers was striking (p < 0.02 for all, p < 0.005 for each alone; Fisher’s exact test). We hypothesized that the increased copy-number in these samples may be due to large-scale amplifications involving the Cyclin E1 (*CCNE1*), a known driver in endometrial, breast and ovarian cancers located on chromosome 19, along with *BRD4*. Indeed, the median copy-number at *CCNE1* was significantly higher in *BRD4*-deletion samples compared to samples without *BRD4* deletions (7.0 vs. 3.0; p < 0.0002; Wilcoxon-rank test; **Figure 1c**). Additionally, tumors with amplicons containing the *BRD4* locus (*BRD4-*amplified) additionally had significantly more concomitant focal deletions in *BRD4* compared to those without gene amplification (p=0.00004) (**Figure 1D**). In each tumor type, areas of genomic deletion overlapped with epigenetic marks of transcriptionally active chromatin, including H3K27ac peaks from ENCODE cell lines (n=7) and H3K4me3 peaks from ovarian (HMEC) and breast adenocarcinoma (MCF-7) cell lines^13^ (**Figure 1A**). We therefore hypothesized that focal *BRD4* deletions would lead to decreased expression levels of *BRD4*, either through copy-number dosage effects or to disruption of key regulatory regions.

### Focal deletions of *BRD4* impact gene isoform expression ratios

We next performed differential expression analysis to identify the genes most dysregulated in tumors with *BRD4* focal deletions. Controlling for tumor type and restricting to autosomal protein-coding genes, we identified 23 genes (of 18,061 genes) with significantly differential expression between the *BRD4* focal deletion and non-deleted cohorts (q < 0.1 cutoff with Bonferroni correction and at least 2-fold expression change; **Figure 2A**). Of these 23 genes, 17 were over-expressed in the *BRD4* focal deletion group. Interestingly, 76% (13/17) of over-expressed genes were located on chromosome 19, including *CCNE1* (**Figure 2B**). *BRD4* did not achieve significance, even in a larger list of 584 genes that achieved significance when performing less restrictive false discovery rate (FDR) correction. We also performed differential sequence analysis without controlling for tumor type, finding *CCNE1* as the single most significant gene in this analysis and with 57% (17/30) of the top 30 genes being found on chromosome 19. Again, there was no significant difference in *BRD4* expression. We thus reasoned that *BRD4* focal deletions prevent toxic *BRD4* over-expression that would have otherwise occurred in the background of chromosome 19 amplifications driving *CCNE1*.

**Figure 2.**
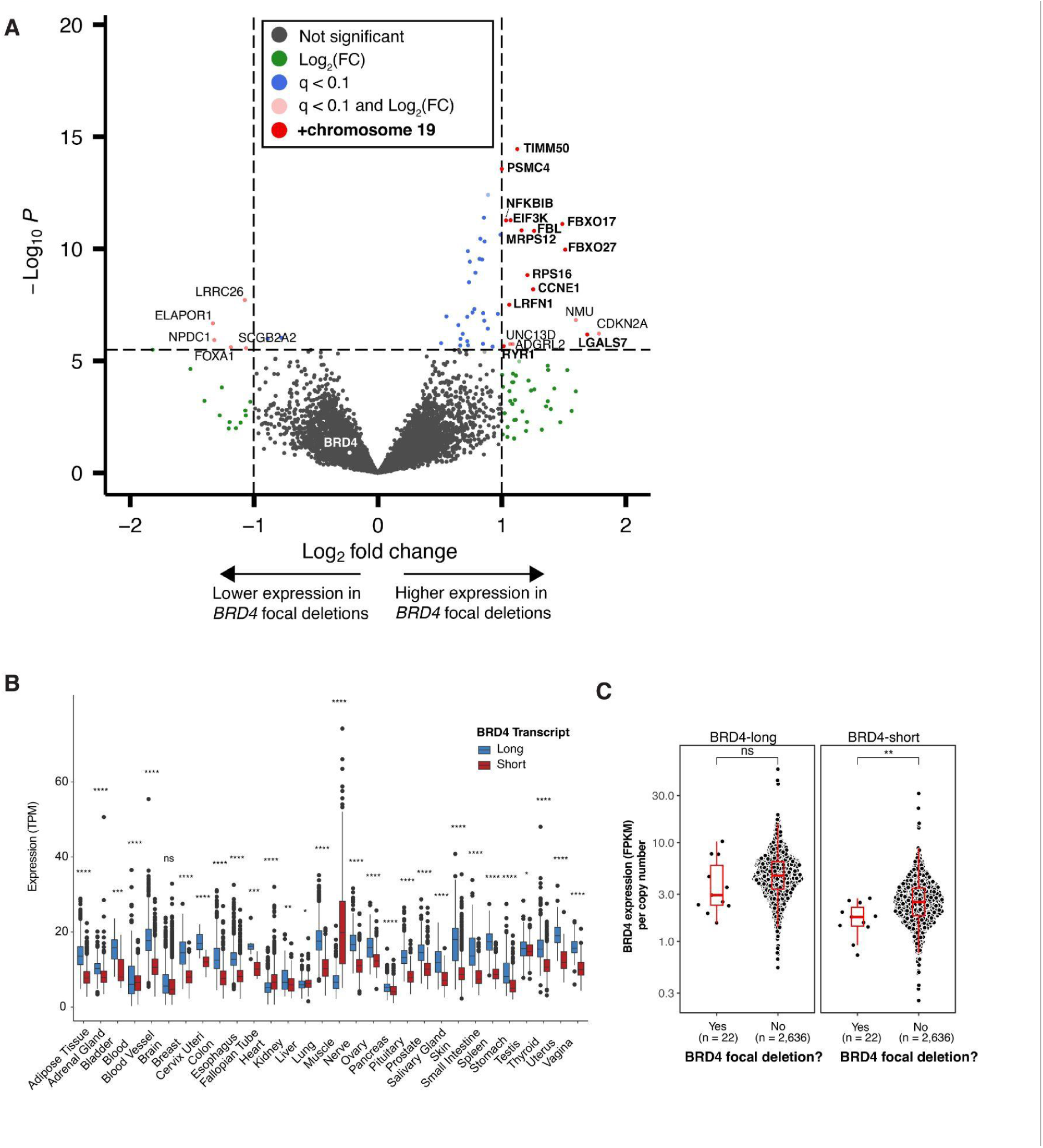
Effect of *BRD4* focal deletions on isoform-specific gene expression. **(A)** Volcano plot of global gene expression comparing *BRD4* focal deletion tumors (n=22) with all other PCAWG tumors (n=2,636), Bonferroni-corrected cutoff of p < 0.1. Genes on chromosome 19 are displayed in bold type. **(B)** Expression of BRD4-long (ENST00000263377) and BRD4-short (ENST00000371835) isoforms in normal tissues as obtained from GTEx (TPM = transcripts per million). Two-tailed t-test. **(C)** Isoform-specific expression of BRD4 in PCAWG data comparing BRD4-long (left) with BRD4-short (right) between tumors with or without *BRD4* focal deletions (FPKM = fragments per kilobase of transcript per million fragments mapped). Wilcoxon rank-sum test.

*BRD4* encodes two splice variants, “long” and “short”, with distinct cellular localization and biological roles. The “long” isoform (BRD4-long) consists of 20 exons and encodes a 1362 amino-acid protein mostly confined to the nuclear membrane. It has both a proline-rich region and a C-terminal motif that allows for interaction with the p-TEFb complex and RNA polymerase II^14^. In contrast, the “short” isoform (BRD4-short) consists of the first 12 exons, encoding a 722 amino-acid protein located in the nuclear matrix. While lacking the proline-rich domain, this isoform modulates chromatin organization and recruitment of transcription factors^15^. In cancer, BRD4-long has been attributed a tumor suppressive role, while BRD4-short is deemed oncogenic and reported to be highly expressed in metastatic disease^16,17^. We therefore asked whether the deletions’ effects on expression favored one of these two isoforms.

We first assessed the distribution of BRD4-long and BRD4-short isoforms across normal tissues, using the Adult Genotype Tissue Expression (GTEx) Project (**Figure 2C**). Except for muscle, BRD4-long is expressed at significantly higher levels (p < 0.0001) than BRD4-short. Given the direct implications for oncogenesis, we conducted analyses in the PCAWG samples to determine whether focal deletions occurring on *BRD4*’s regulatory regions could impact, in addition to global gene expression, the ratio of expressed isoforms (**Figure 2**). When normalized for locus-specific copy-number, we identified significantly decreased copy-number adjusted expression of the BRD4-short isoform (p = 0.017) in tumors with *BRD4* focal deletions. There was no significant difference in copy-number adjusted BRD4-long expression (p = 0.11). Taken together, these data provide evidence of how rearrangements may alter gene isoform ratios and ultimately reduce over-expression of a gene residing in a genomically amplified region.

**Extended Figure 1.**
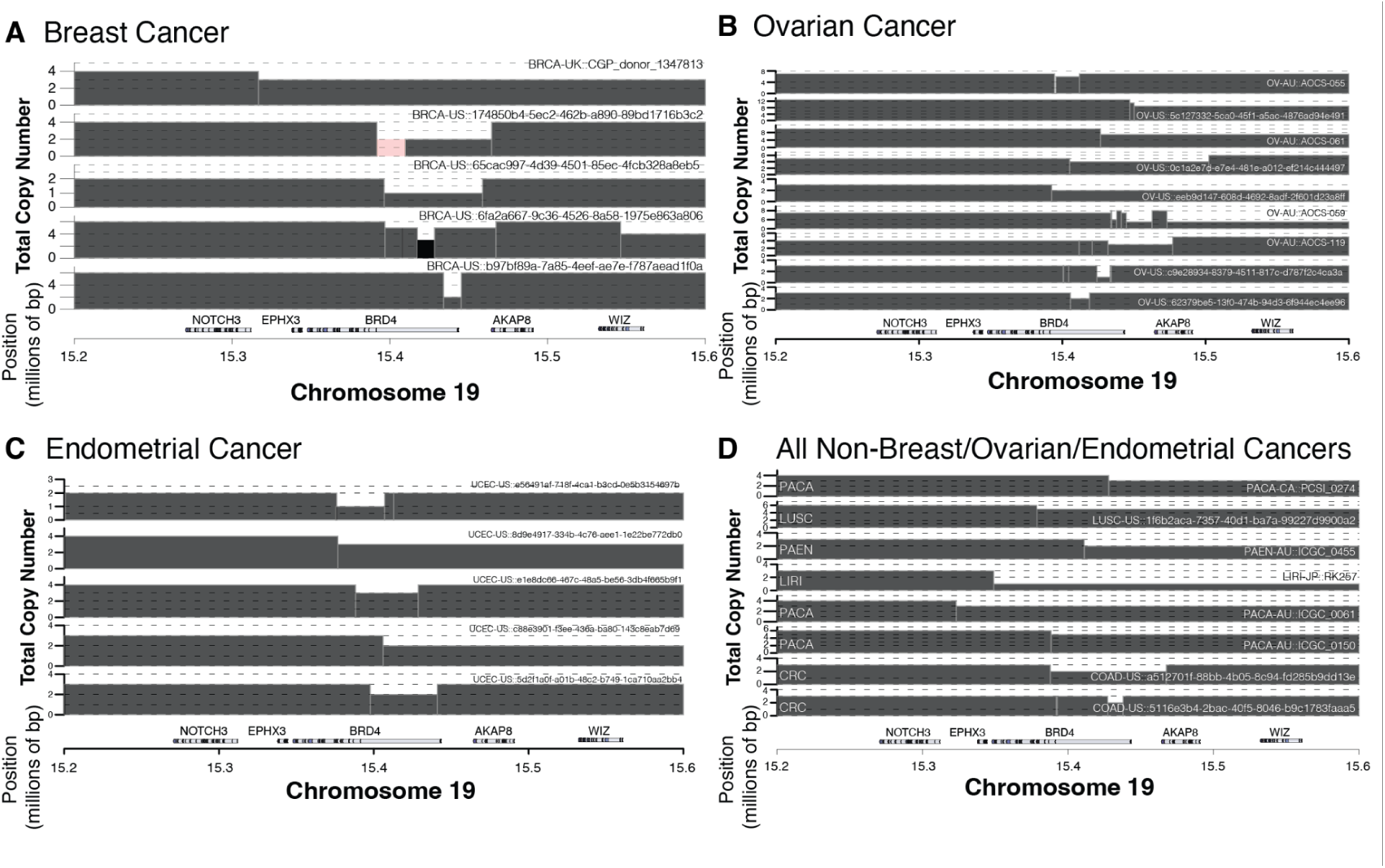
SCNA events at the *BRD4* locus corresponding to the recurrent breakpoints. Total copy-number for tumors with either a focal deletion at the *BRD4* locus or an SCNA breakpoint within the *BRD4* locus for **(A)** breast cancer (n=5) **(B)** ovarian cancer (n=9) **(C)** endometrial cancer (n=5) and **(D)** all other cancers including pancreatic adenocarcinoma (PACA; n=3), pancreatic neuroendocrine tumor (PAEN; n=1), lung squamous cell carcinoma (LUSC; n=1), hepatocellular carcinoma (LIRI; n=1) and colorectal cancer (CRC; n=2). Only those tumors with *BRD4* focal deletions are included in subsequent analyses.

**Extended Figure 2.**
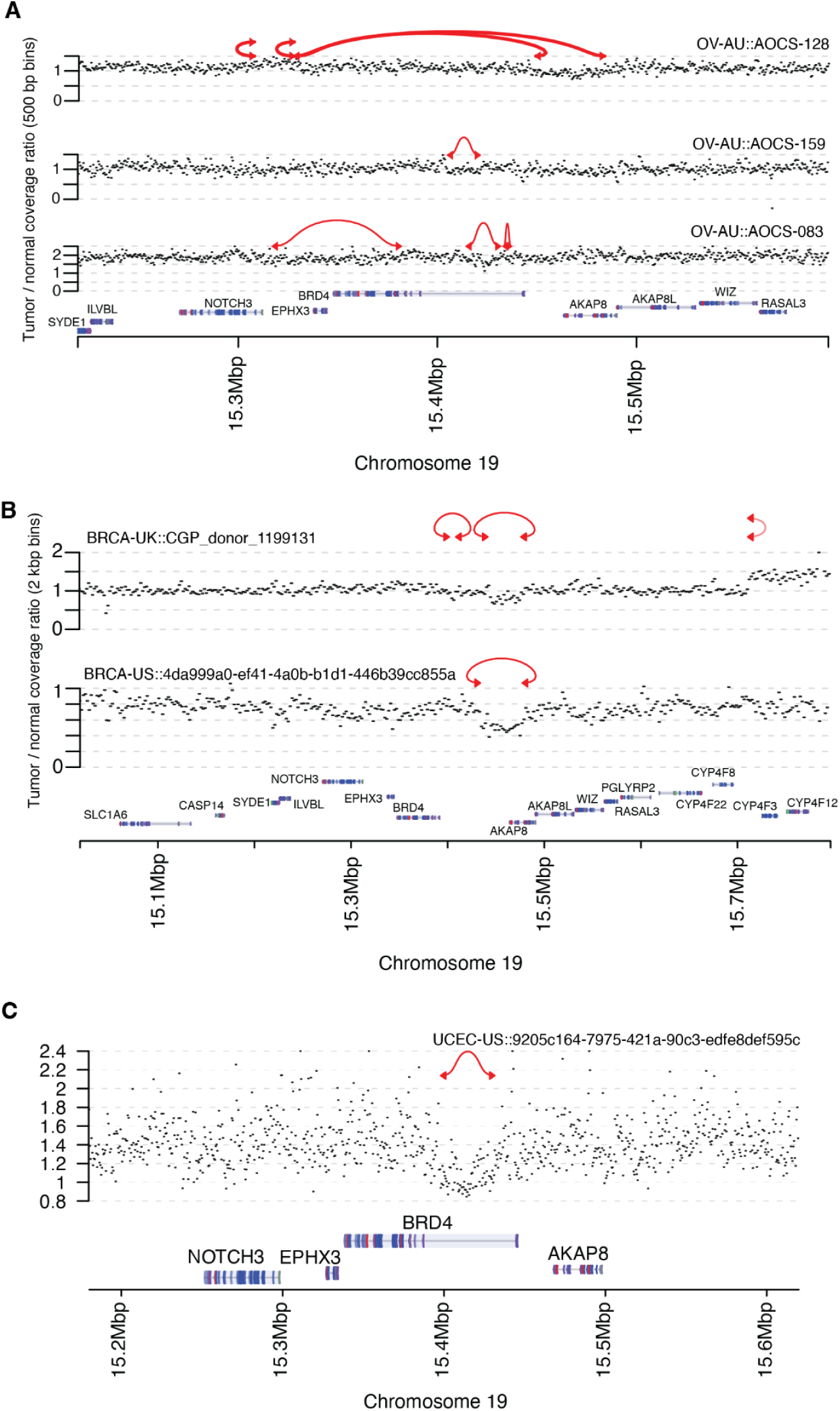
Tumors with breakpoints within the recurrent region of *BRD4*, but with no corresponding SCNAs. Tumor/normal coverage ratios from binned read-depth signals (y-axis) and rearrangements (red lines; arrows indicate breakpoint orientation) for tumors without a called SCNA from read-depth only signal for **(A)** ovarian cancer (n=3), **(B)** breast cancer (n=2), and **(C)** endometrial cancer (n=1).

### BRD4 gene overexpression is detrimental for cancer cell proliferation

We hypothesize that *BRD4* is initially amplified in cancer cells, but high expression levels are not tolerated. *BRD4* gene over-expression has been previously linked with a growth inhibition phenotype in HeLa cervical cancer cells^19^, but the effects of gene overexpression across multiple cancer types has not been systematically interrogated. We therefore compiled data from the “control” arm of 15 different open reading frame (ORF) overexpression screens performed in 8 tumor types^20–28^, where the effects of gene overexpression on cell growth can be quantified over time. We next determined the log-fold changes in cell growth following overexpression of BRD4-long and BRD4-short isoforms across different cell lines for a period of 2-3 weeks and in the absence of any additional selection pressures **(Figure 3A**). In 10/16 cell lines (62.5%), global *BRD4* overexpression decreased cell growth (as evidenced by negative log-fold changes), supporting its role as a “toxic” gene in cancer^29^. For the remaining cell lines, gene overexpression had a neutral effect on cell proliferation (**Figure 3B**). Within each cell line, overexpression of BRD4-short was significantly more toxic (p = 0.001) than BRD4-long (**Figure 3C**).

**Figure 3.**
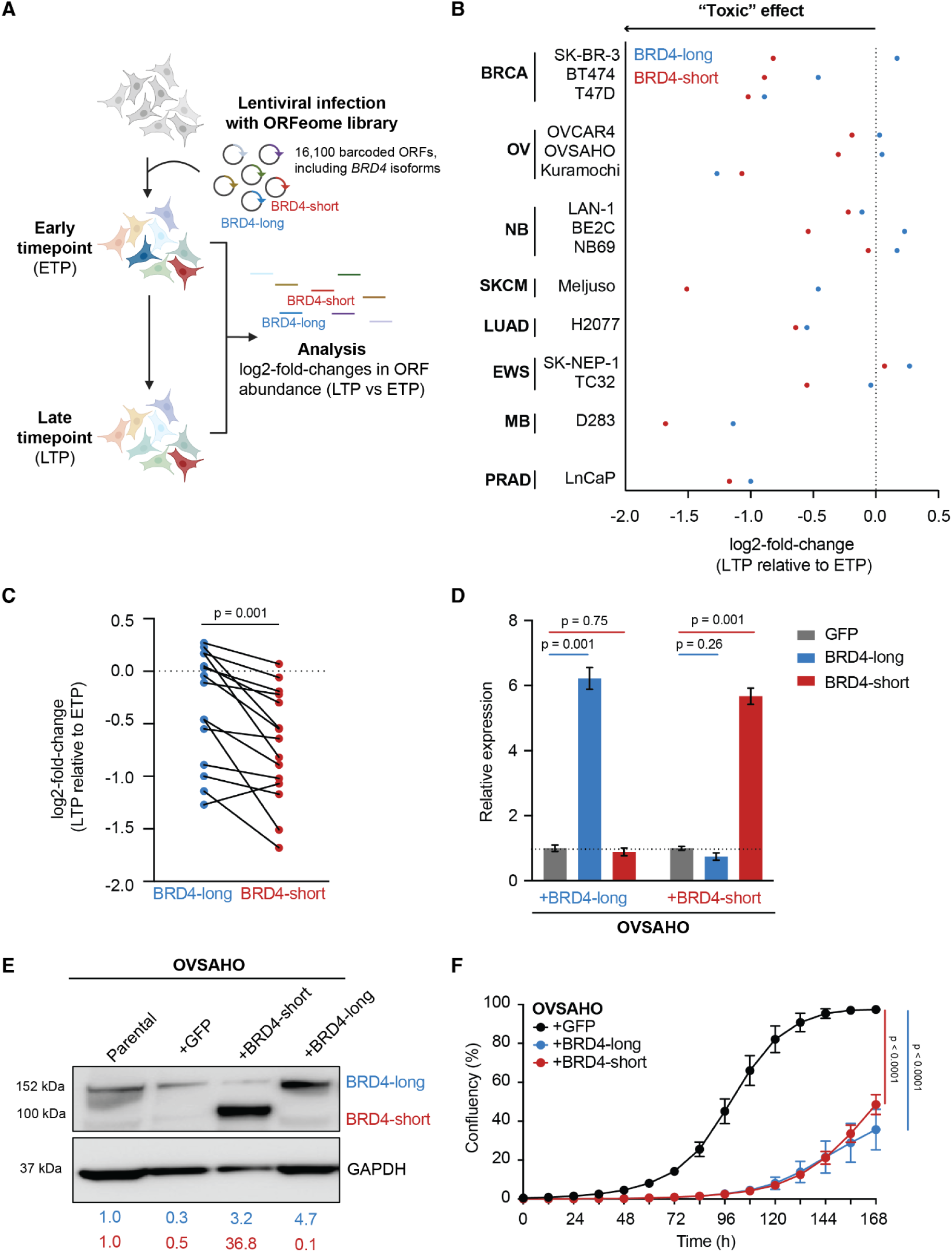
*BRD4* overexpression is toxic across multiple cancer types. **(A)** Overview of an ORF screen, from which the effect of BRD4-long and BRD4-short overexpression on cell proliferation can be assessed. **(B)** Log_2_-fold-changes in cell proliferation for BRD4 constructs, expressed relative to the experimental early time point (ETP) of each screen. Negative changes reflect a detrimental effect on cell proliferation. ORF screens were conducted in breast (BRCA), ovarian (OV), neuroblastoma (NB), skin cutaneous melanoma (SKCM), lung adenocarcinoma (LUAD), Ewing sarcoma (EWS), medulloblastoma (MB) and prostate adenocarcinoma (PRAD) cell lines. **(C)** ORF screen performance between BRD4-long and BRD4-short isoforms for each cell line. Two-tailed t-test. **(D)** Validation of BRD4 isoform expression in OVSAHO cells. Transcript expression is quantified relative to GFP control. Mean and standard deviation of three replicates, two-way ANOVA. **(E)** Immunoblot of BRD4-long and BRD4-short protein variants detected in OVSAHO cells transduced with BRD4 or control GFP overexpression constructs. Values represent levels of protein expression, normalized to GAPDH expression and shown relative to parental cells (endogenous BRD4 levels). **(F)** Quantification of cell confluency in OVSAHO cells transduced with BRD4-long, BRD4-short or GFP control overexpressing constructs. Mean and standard error of three replicates, two-way ANOVA.

We further selected OVSAHO ovarian carcinoma cells as a model to validate the toxic effects associated with *BRD4* overexpression from the ORF screen data. We focused on ovarian tumors as they present higher *BRD4* gene amplification rates (∼12%) when compared to breast cancers (∼3%)^30^. First, we stably transduced OVSAHO cells with a lentiviral vector encoding BRD4-long, BRD4-short, or GFP control. Following cell selection, we validated gene overexpression by RT-qPCR using isoform-specific primers. We were able to detect >5-fold increases in BRD4-long and BRD4-short transcripts (p < 0.001) when compared to GFP-transduced OVSAHO cells (**Figure 3D**). These changes translated into BRD4 protein expression levels, where BRD4-long and BRD4-short isoforms were expressed 4.6-fold and 36.8-fold, respectively, compared to GFP controls (**Figure 3E**). To assess the effects of *BRD4* over-expression on cell proliferation, we conducted live-cell image analysis to quantify increases in confluency present in cells over-expressing different isoforms over the course of 7 days (**Figure 3F**). Over-expression of both BRD4-long and BRD4-short led to a profound growth inhibitory phenotype (p < 0.001) compared to GFP over-expressing cells, confirming that supraphysiological *BRD4* gene expression levels have a negative impact on cellular fitness.

### *BRD4* focal deletions rescue ovarian carcinoma cells from toxicity effects associated with gene overexpression

The presence of focal deletions spanning *BRD4* regulatory regions suggests that these rearrangements may mechanistically reduce gene expression to levels that are no longer toxic to cancer cells. To functionally validate this hypothesis, we first stably infected OVSAHO cells with a lentiviral Cas9 vector and confirmed protein expression (**Figure 4A**) and activity (**Figure 4B**) 10 days following selection. Next, we transduced cells with CRISPR-Cas9 single-guide RNAs (sgRNAs) designed to target two intronic regions (intron 1 of RefSeq^18^ transcript BRD-004) of *BRD4* (“sgBRD4_region1” and “sgBRD4_region2”), mimicking the effect of the focal deletions originally identified in PCAWG samples (**Figure 4C**). We were able to confirm sgRNA cutting by CRISPR-sequencing (**Figure 4D**), detecting 46.4% of frameshift cuts for sgBRD4_region1 and 66.4% for sgBRD4_region2 when amplicon reads were aligned to a reference *BRD4* sequence. To assess global and isoform-specific levels of *BRD4* expression following sgRNA cutting, we performed a RT-qPCR in OVSAHO-Cas9 cells transduced with *BRD4* sgRNAs relative to a sgGFP control (**Figure 4E**). Global *BRD4* levels were decreased by ∼40% following sgRNA cutting (p < 0.0001). No difference in the ratio of long to short isoforms was detected when regions 1 or 2 of *BRD4* were deleted (p > 0.99). As deletion of these intronic or exon *BRD4* regions decrease gene expression levels, we next evaluated whether they would affect cell proliferation. By live-cell imaging, we were able to confirm that deletions by sgBRD4_region1 and sgBRD4_region2 significantly decrease cell confluency by ∼50% at 7 days compared to cells transduced with a sgGFP control (p < 0.0001) (**Figure 4F**). The observation that *BRD4* ablation is detrimental to OVSAHO cell growth is in agreement with CRISPR-Cas9 dependency data from the Dependency Map Consortium (DepMap Public 23Q4 release; http://depmap.org), where *BRD4* scores as a genetic dependency (mean Chronos score = -0.99 ± 0.28) and is essential in 33/59 (∼56%) of ovarian cancer cell lines (**Figure 4G**).

**Figure 4.**
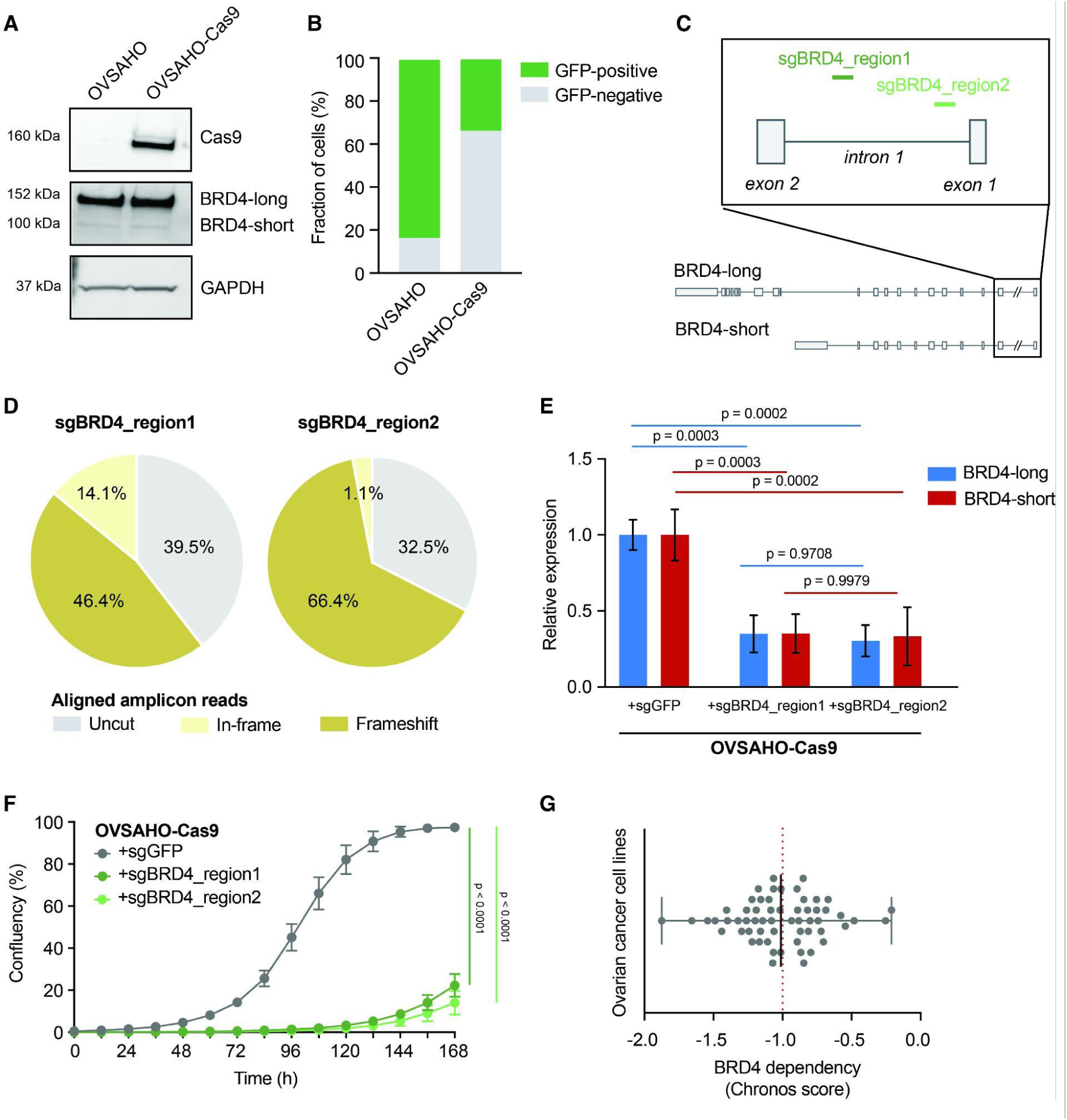
Use of CRISPR-Cas9 technology to mimic the effect of *BRD4* focal deletions. **(A)** Confirmation of Cas9 expression in OVSAHO cells by immunoblotting. **(B)** Fraction of GFP-positive cells detected in Cas9 activity assay. To measure Cas9 cutting efficiency, OVSAHO-Cas9 cells were transduced with a vector encoding GFP and sgGFP. After 10 days, a decreased population of GFP-positive cells was detected by flow cytometry, confirming Cas9 activity. **(C)** Intronic *BRD4* regions targeted by CRISPR-Cas9 sgRNAs (inset not to scale due to length of intron 1). **(D)** CRISPR-sequencing results for sgBRD4_region1 and sgBRD4_region2, showing the percentage of uncut, frameshift, and in-frame *BRD4* reads when aligned to a reference sequence. **(E)** Expression of *BRD4* isoforms following CRISPR-Cas9-mediated sgRNA cutting, relative to sgGFP control. Median and standard deviations of three replicates, two-way ANOVA **(F)** Quantification of cell confluency in OVSAHO-Cas9 cells following CRISPR-Cas9-mediated sgRNA cutting of *BRD4* regulatory regions or sgGFP control. Mean and standard error of three replicates, two-way ANOVA. **(G)** Landscape of *BRD4* dependency across ovarian cancers. Chronos scores (with mean and standard deviation) are shown for every ovarian cancer cell line included in the DepMap Public 23Q4 release. A score of -1 (median score for essential genes; red dashed line) is used as reference to indicate genetic dependency.

We finally attempted to functionally recreate the gene regulation patterns observed in *BRD4*-amplified PCAWG tumors that undergo focal rearrangements at the *BRD4* locus. This would allow us to confirm that the observed focal deletions indeed reduce and finetune levels of *BRD4* expression to rescue cells from gene toxicity and ultimately sustain proliferation. To “time” *BRD4* overexpression with *BRD4* deletion, we first transduced OVSAHO Cas9-expressing cells with sgBRD4_region1 and sgBRD4_region2, and later introduced BRD4-long or BRD4-short according to previously established experimental timepoints (**Figure 5A**). After 7 days, we observed that deletion of sgBRD4_region1 and sgBRD4_region2 rescued proliferation of BRD4-long and BRD4-short OVSAHO cells compared to a sgGFP control (**Figure 5B**). Altogether, our results point to a ‘Goldilocks’ model of *BRD4* gene regulation, where too high or too low levels of expression have a negative effect on cellular fitness. In *BRD4-*amplified tumors, focal deletions in gene regulatory regions serve as a mechanism to decrease and restore gene expression to levels that are tolerated and beneficial for proliferation (**Figure 5C**). This represents a novel mechanism of gene-dosage compensation in human cancers.

**Figure 5.**
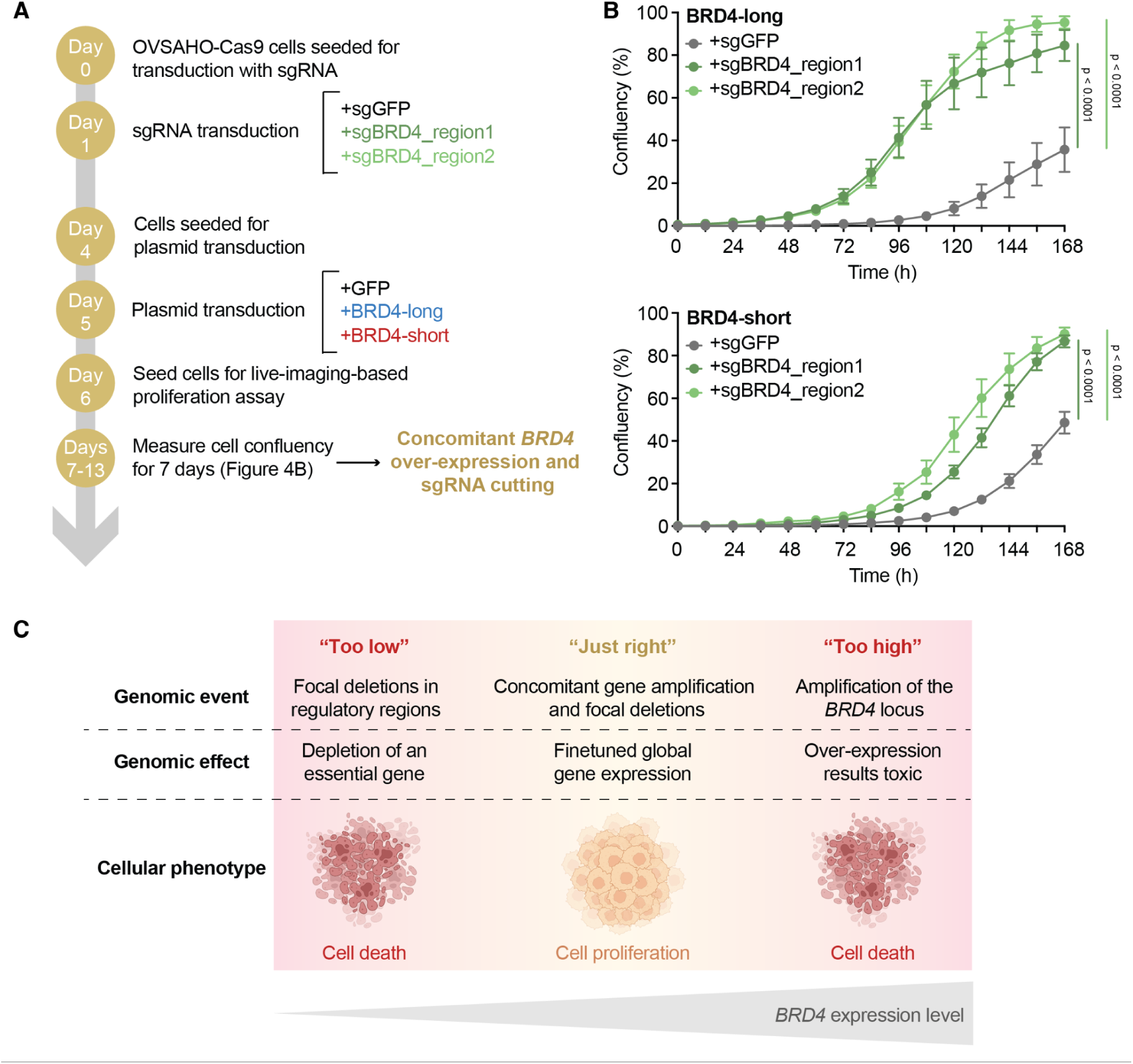
Finetuned levels of *BRD4* global expression are required for sustained cellular proliferation. **(A)** Experimental workflow for mimicking concomitant focal *BRD4* deletions occurring in *BRD4* over-expressing ovarian cancer cells. **(B)** Rescue experiment by modulation of *BRD4* expression. Quantification of cell confluency after CRISPR-Cas9-mediated gene ablation was measured in OVSAHO-Cas9 cells over-expressing BRD4-long (top) or BRD4-short isoforms (bottom). Mean and standard error from three replicates, two-way ANOVA. **(C)** Model of *BRD4* expression regulation. In ovarian cancers, too low or too high expression levels are detrimental to cellular fitness. In *BRD4*-amplified tumors, focal deletions serve as a mechanism to dampen global gene expression and rescue cellular proliferation.

## Discussion

The distribution of rearrangements across the cancer genome not only reflects the mechanisms that give rise to their formation (e.g. homologous recombination, microhomology-dependent recombination, retrotransposition), but also the advantage they confer for cellular fitness^31,32^. PCAWG’s initial analysis of non-coding somatic drivers in over 2,600 cancer whole genomes identified breast and ovarian tumors with recurrent breakpoints in the *BRD4* locus on chromosome 19p, which result in focal deletions that span gene regulatory regions and reduce global expression levels^3^. Our present study revealed that these structural variations further affect the ratio of expressed *BRD4* isoforms, and provided functional evidence for these deletions serving as a mechanism to finetune gene expression.

*BRD4* overexpression has been previously linked with the suppression of cancer cell growth in selected human and mouse cell line models^19^, but has not been attributed with the toxicity profile we observed when systematically interrogating the effects of gene overexpression across multiple tumor types. These toxicity patterns seem to additionally differ between BRD4-long and BRD4-short isoforms. This suggests the importance of characterizing which *BRD4* variants become deregulated in a particular cancer type, as they exert different oncogenic roles and may inform the most appropriate strategies for targeted therapy. Functional evidence from our ORF screen analysis further highlights the biological relevance of *BRD4* deregulation in other cancer types, such as medulloblastoma, where targeted therapies against BRD4, such as BET-bromodomain inhibitors, hold clinical promise^23,33^.

This is not the first case where genomic amplifications are toxic to cancer. In recent efforts, our group and others have characterized the effect of collateral amplifications occurring across multiple tumor types, identifying genes that cause proliferation defects when overexpressed due to their close genomic proximity to an amplified oncogene. Such amplifications introduce genetic vulnerabilities and may present novel therapeutic avenues^29,34–36^. Our current study provides evidence of *BRD4* focal deletions serving as a mechanism of gene-dosage compensation following gene amplification.

## Methods

### Structural variant analysis

We first analyzed PCAWG samples with amplifications and somatic SV breakpoints overlapping the *BRD4* locus using GISTIC gene-level thresholded copy number data and consensus-called SV and SCNA callsets from the ICGC Data Portal’s DCC Data Release [https://dcc.icgc.org/releases/PCAWG/consensus_cnv/GISTIC_analysis]. We identified cases with focal *BRD4* deletions as those cases with an SCNA breakpoint within (or within 50 Kbp) of the *BRD4* locus leading to a relative copy-loss within *BRD4*, and with a total copy-loss SCNA event size of < 100 Kbp. An additional 6 cases of *BRD4* focal deletions were identified by manual review of tumor/normal read depth plots, with SVs (rearrangements) annotated, for all breast, ovarian and endometrial cancers containing *BRD4* breakpoints. These cases were deemed to have a true focal deletion if there was a visible decrease in read-depth signal with drop-offs corresponding to the breakpoints from the SV data, and if those breakpoint orientations were compatible with a copy-loss (outgoing orientation from the deletion) consistent with a deletion.

Total copy-number for any given locus (*e.g. CCNE1*) was taken as the mean total copy-number across that gene, using hg19 genomic coordinates obtained from Ensembl^37^. For calculating copy-number normalized gene expression of *BRD4*, we took the copy-number to be that of the neighboring gene *NOTCH3*, since the copy-number within *BRD4* for *BRD4* focal deletion samples was variable by definition. We performed significance testing using a Wilcoxon rank-sum test with the R stats package (v. 4.2.1). We identified tumors containing an amplification spanning the *BRD4* locus by intersecting the SCNA data for events with total copy-number > 2 with the *BRD4* locus. This was determined irrespective of whether there was a concomittant *BRD4* focal deletion occuring on top of an amplification. We performed a Fisher’s exact test with the R stats package (v. 4.2.1) to determine if *BRD4* focal deletions were enriched in amplified *BRD4* loci, using a p-value significance cutoff of 0.05.

### BRD4 expression analysis

Gene level and transcript level RNA expression data was downloaded from the PCAWG data portal as above. RNA expression values were used as previously described and without further modification^38^. BRD4-long corresponded to Ensembl transcript ENST00000263377. BRD4-short corresponded to Ensembl transcript ENST00000371835. Differential expression analysis was performed using the limma package (v. 3.58.1^39^) in R and the voom method for linear modeling^40^. Expression analysis was restricted to just protein coding genes on chromosomes 1 through 22. We used a model matrix consisting of either (1) just the tumor class (*BRD4* deletion versus not) or (2) tumor class with the tumor type as obtained from the DCC project code (e.g. BRCA tumors consisted of all tumors from projects: BRCA-EU, BRCA-US and BRCA-UK).

For the analysis of *BRD4* isoform expression across normal tissues, we downloaded transcript per million (TPMs) measurements from the GTEx Analysis V7 RNA-sequencing data [https://gtexportal.org/home/downloads/adult-gtex/bulk_tissue_expression], and compared transcript expression levels across available tissue types using paired t-tests with an FDR correction

### Analysis of open reading frame (ORF) screens

The analysis of ORF screen data was performed as previously described^29^. Briefly, we looked at data from 16 independent screens conducted at the Broad Institute of MIT and Harvard, were cells were infected with the ORFeome pLX_317 barcoded library, which contains 16,100 barcoded ORFs over-expressing 12,753 genes. From these, we compared ORF abundance between early and late timepoints for BRD4-long (TRCN0000477053; NM_058243.2) and BRD4-short (TRCN0000491254; NM_014299.2) isoforms. A negative log2-fold change in cell growth following ORF expression as interpreted as an indication of gene toxicity.

### Cell line culture conditions

OVSAHO human high-grade serous ovarian cancer cells were commercially acquired (cat. no. SCC294, Sigma-Aldrich) and cultured in Advanced RPMI 1460 Medium (cat. no. 12633012, ThermoFisher Scientific) supplemented with 10% heat inactivated Fetal Bovine Serum (FBS) and 1% Pen Strep (cat. no. 16140071 and 15070063, ThermoFisher Scientific). Human HEK-293T were commercially acquired (cat. no. CRL-11268, ATCC) and grown in Advanced DMEM (cat. no. 12491015, ThermoFisher Scientific) medium supplemented with 10% FBS. All cell lines were incubated at 37^॰^C with 5% CO_2_, fingerprinted, and frequently checked for mycoplasma contamination using the MycoAlert Mycoplasma Detection Kit (cat. no. LT07-318, Lonza).

### Generation of lentiviral overexpression constructs

pDONR223 vectors encoding the BRD4-long (clone ID: ccsbBroadEn_11738; NM_058243.3) and BRD4-short (clone ID: ccsbBroadEn_11738; NM_014299.2) isoform sequences, or a GFP control (clone ID: BRDN0000464762), were acquired from the Genomic Perturbation Platform (GPP) at the Broad Institute of MIT and Harvard. Next, each coding sequence was cloned into the pLX_307 lentiviral vector (GPP, Broad Institute of MIT and Harvard), driven by a EF1a promoter and under puromycin selection, using the Gateway LR Clonase II enzyme mix (cat. no. 12535-029, Invitrogen) according to the manufacturer’s protocol. The resulting products from the LR recombination reaction were then transformed into One Shot Stabl3 Chemically Competent *E. coli* (cat. no. C737303, ThermoFisher Scientific) at 37^॰^C in LB microbial growth medium containing 100 ug/mL carbenicillin. Plasmid DNA was extracted from single colonies using QIAprep Spin Miniprep (cat. no. 27104, Qiagen) and Plasmid Plus Maxi Kits (cat. no. 12963, Qiagen). Lentiviral transduction was performed by co-transfection of HEK-293T cells with 10 μg of pLX_307 vector (BRD4 or GFP control), 1 μg psPAX2 (plasmid #12260, Addgene) and 1 μg pVSV-G (plasmid #138479, Addgene) viral packaging plasmids using Lipofectamine 3000 Transfection Reagent (cat. no. L3000001, ThermoFisher Scientific). After 48 h, virus harvesting was conducted using a 0.45 μm syringe. For cell transduction, 400 μL of virus were added to OVSAHO cells seeded at a density of 2 million cells per well in a 12-well plate. After an overnight incubation at 37^॰^C, cells were selected for 5 days with 1 µg/mL puromycin (cat. no. A1113803, ThermoFisher Scientific).

### Generation of lentiviral Cas9 and single-guide RNA constructs

The lentiviral vectors pXPR_BRD111 and pXPR_BRD016 expressing human SpCas9 and a single-guide RNA (sgRNA) of interest, respectively, were acquired from the GPP Platform (Broad Institute of MIT and Harvard). While SpCas9 lentiviral expression is under blasticidin selection, expression of the sgRNA of interest is driven by a human U6 promoter in a construct under hygromycin selection. The sgRNAs against BRD4_region1 and BRD4_region2 were cloned into the pXPR_BRD016 backbone following the lentiCRISPRv2 one vector system protocol described by the Zhang laboratory^41,42^. The oligos designed against sgBRD4_region1 were: forward 5’-CACCGTCTACCACACTGTAATGCTG-3’, and reverse 5’-AAACCAGCATTACAGTGTGGTAGAC-3’. The oligos designed against sgBRD4_region2 were: forward 5’-CACCGCGTAGTCTTGAGTAAAAGGG-3’, and reverse 5’-AAACCCCTTTTACTCAAGACTACGC-3’. As a control, we cloned a sgRNA targeting GFP (“sgGFP”) using oligos forward 5’-CACCGGAGCTGGACGGCGACGTAAA-3’ and reverse 5’-AAACTTTACGTCGCCGTCCAGCTCA-3’. Lentiviral production and transduction were performed in HEK-293T and OVSAHO cells as described above. Selection of OVSAHO Cas9 over-expressing cells was conducted for 7 days with 10 µg/mL blasticidin (cat. no. A1113903, ThermoFisher Scientific). After transduction with sgRNA-expressing constructs, a second round of selection was performed for 10 days with 100 µg/mL hygromycin B (cat. no. 10687010, ThermoFisher Scientific).

### Quantification of gene expression by quantitative reverse transcription PCR (RT-qPCR)

RNA was first extracted from one million cells using the RNeasy Mini Kit (cat. no. 74004, Qiagen) using an on-column DNase digestion (cat. no. 79254, Qiagen). Next, cDNA conversion was conducted from 2 µg purified RNA using the Maxima H Minus first-strand cDNA synthesis kit (cat. no. K1652, ThermoFisher Scientific). Quantitative PCR analysis was done on 500 ng cDNA mixed with 1X Maxima SYBR Green/ROX qPCR Master Mix (cat. no. K0221, ThermoFisher Scientific) and 0.3 µM primers. For the detection of *BRD4* isoform transcripts, the following primers were used: BRD4-long forward 5’-CCGGCTCCTCCAAGATGAAG-’3’ and reverse 5’-ATGGTGCTTCTTCTGCTCCC-3’; BRD4-short forward 5’-CCTCAAGCTGAGAAAGTTGATGTG-’3’ and reverse 5’-AGGACCTGTTTCGGAGTCTT-3’. As an endogenous control, primers forward 5’-CCGAAAGTTGCCTTTTATGG-3’ and reverse 5’-TCATCATCCATGGTGAGCTG-3’ targeting *ACTB* were used. Samples were run in a StepOne thermocycler according to the following protocol: 40 cycles of 95°C for 15 sec and 60°C for 1 min, melt curve stage at 95°C for 15 sec, and 60°C for 1 min. Quantification of mRNA expression relative to control was performed by the ΔΔCT method.

### Immunoblotting

Two million cells were pelleted and lysed in RIPA buffer, containing 20 mM Tris-HCl at pH 7.5, 150 nM NaCl, 1 mM Na_2_EDTA, 1 mM EGTA, 1% NP-40, 1% sodium deoxycholate, 2.5 mM sodium pyrophosphate, 1 mM beta-glycerophosphate, 1 mM Na_3_VO_4_, and 1 μg/mL leupeptin. After a 30 min incubation on ice, samples were centrifuged at 13,000 rpm for 10 min at 4°C. Protein concentrations were quantified using Pierce BCA Protein Assay (cat. no. 23225, ThermoFisher Scientific). Next, 30 µg of each sample was loaded on a NuPAGE 4-12% Bis-Tris protein gel (cat. no. NP0326, ThermoFisher Scientific) and separated using NuPAGE MOPS Running Buffer (cat.no. NP0001, ThermoFisher Scientific) for 2 h at 120 V. A dry gel transfer was completed at 30 V for 6 min using PVDF Transfer Stacks (cat. no. IB24002, ThermoFisher Scientific) in an iBlot 2 instrument (Invitrogen). After blocking with 5% dry milk in TBS-T for 30 min, the membrane was incubated at 4°C overnight with primary antibodies against rabbit BRD4 (cat. no. A301-985A100, ThermoFisher Scientific), mouse SpCas9 (cat. no. 14697, Cell Signaling Technology), or rabbit GAPDH (cat. No. 2118, Cell Signaling Technology). Next, the membrane was washed in TBS-T and incubated for 1h with goat anti-rabbit (cat. no. 7074S, Cell Signaling Technology) or goat anti-mouse (cat. no. 7076S, Cell Signaling Technology) secondary antibodies. Chemiluminescent signal was detected using SuperSignal West Pico PLUS Substrate (cat. no. 34580, ThermoFisher Scientific) in an ImageQuant LAS 4000 imager (GE Healthcare Life Sciences).

### Cas9 activity assay

One million OVSAHO or OVSAHO-Cas9 expressing cells were transduced with the lentivirus pXPR_011 (GPP, Broad Institute of MIT and Harvard), which co-expresses a sgRNA against EGFP and EGFP as a target under puromycin selection. Following infection with 480 µL virus, cells were spun for 2 h at 1,000 *g* at 30°C, and cell media was replaced after 6 h. The next day, 1 µg/mL puromycin was added. To quantify Cas9 activity, the percentage of GFP-negative cells (where Cas9-mediated cutting of EGFP has occurred) was quantified in OVSAHO and OVSAHO-Cas9 cells by flow cytometry using a BD LSRFortessa Cell Analyzer (BD Biosciences) using a 488 nm laser. Non-expressing Cas9 OVSAHO cells appear green and provide a baseline reference for determining Cas9 activity in OVSAHO-Cas9 cells.

### CRISPR amplicon sequencing

To confirm CRISPR-mediated cutting of sgBRD4_region1 and sgBRD4_region2, we first designed primers spanning the regions of interest in *BRD4*’s intron 1. For “region 1”, we used forward 5’-AGATCCTTTTGGCTCCCTGT-3’ and reverse 5’-GGGAGAGAGAGGTTGCAGTG-3’ primers. For “region 2” we used forward 5’-TTGCTTGAAGATGGGAAACC-3’ and reverse 5’-ACGGTAGGGAACTTGACAGC-3’ primers. Next, we amplified each region by PCR using 1× Phusion HF Buffer (Thermo Scientific), 0.2 mM dNTPs, 0.2 μM forward, and 0.2 μM reverse primers, 0.02 U μL−1 Phusion DNA Polymerase (Thermo Scientific) and 6 ng of genomic DNA. The PCR was performed in a 2720 Thermal Cycler (Applied Biosystems) following the protocol: 95°C for 30 sec, 30 cycles of i) 95°C for 10 sec, ii) 58°C for 15 sec, and iii) 72°C for 20 sec, and 72°C for 10 min. CRISPR amplicon sequencing was performed by next-generation sequencing at the CCIB DNA Core (Massachusetts General Hospital). All reads were aligned to NCBI’s *BRD4*’s reference sequence and analyzed with CRISPResso2^43^ to obtain editing frequencies and indel size distribution for each targeted region.

### Proliferation curves

Cells were seeded at a density of 2,000 cells per well in 96-well plates. The next day, live-cell imaging was conducted in an Incucyte Live-Cell Analysis system (Sartorius) and images were captured for each well every 12 h with a 10x objective. Custom-defined masks were then applied to each image to identify cell area and determine confluency as a surrogate of cell growth.

### Analysis of BRD4 dependency

CRISPR dependency scores (Chronos) were downloaded from the DepMap Public 23Q4 release for all available ovarian cancer cell lines (n=59). A Chronos score threshold of -1 was used to call *BRD4* a genetic dependency in a given cell line.

