## Supplementary figures and images for "Recurrent breakpoints in the *BRD4* locus reduce toxicity associated with gene amplification"

### Extended Figure 1

*Extended Figure 1*

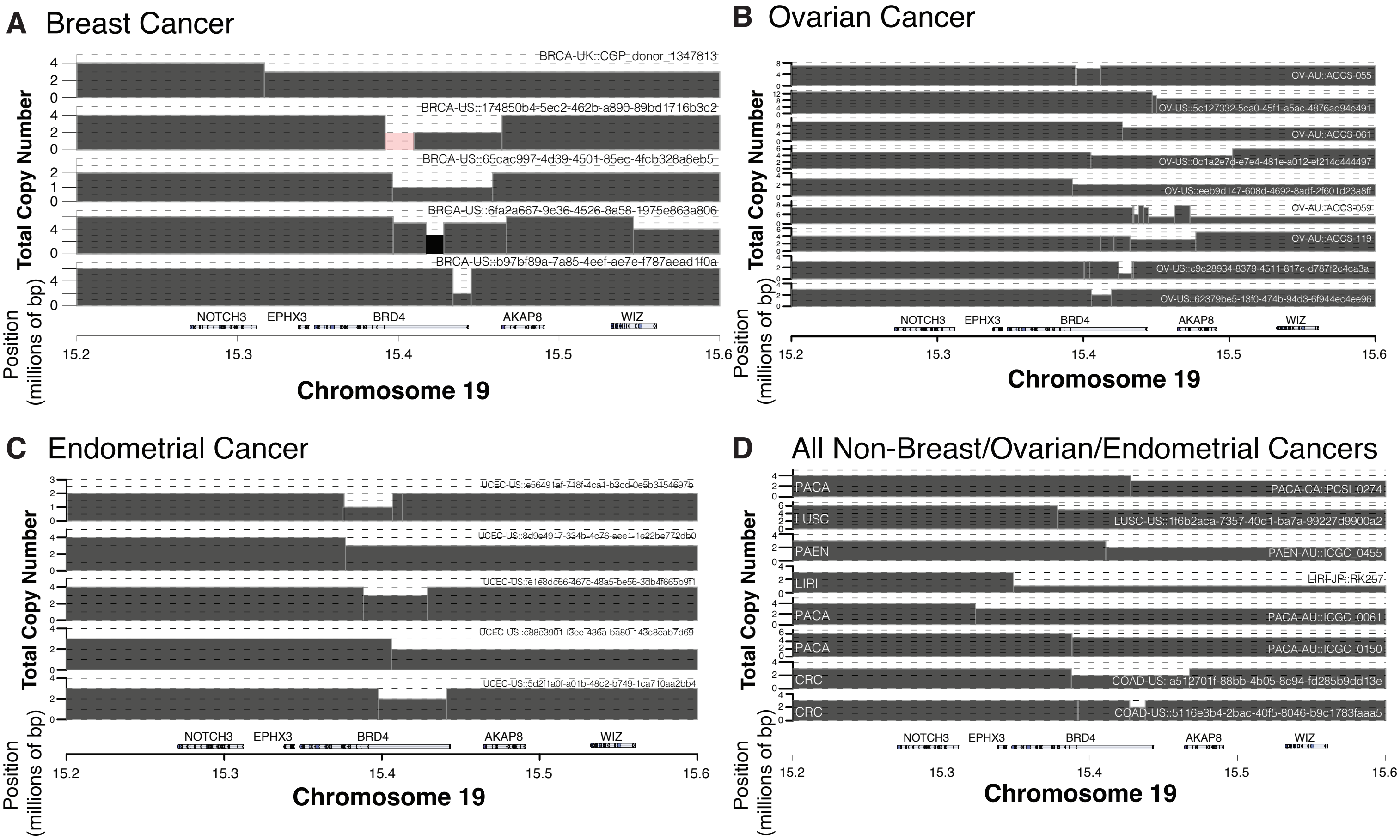

### Extended Figure 2

*Extended Figure 2*

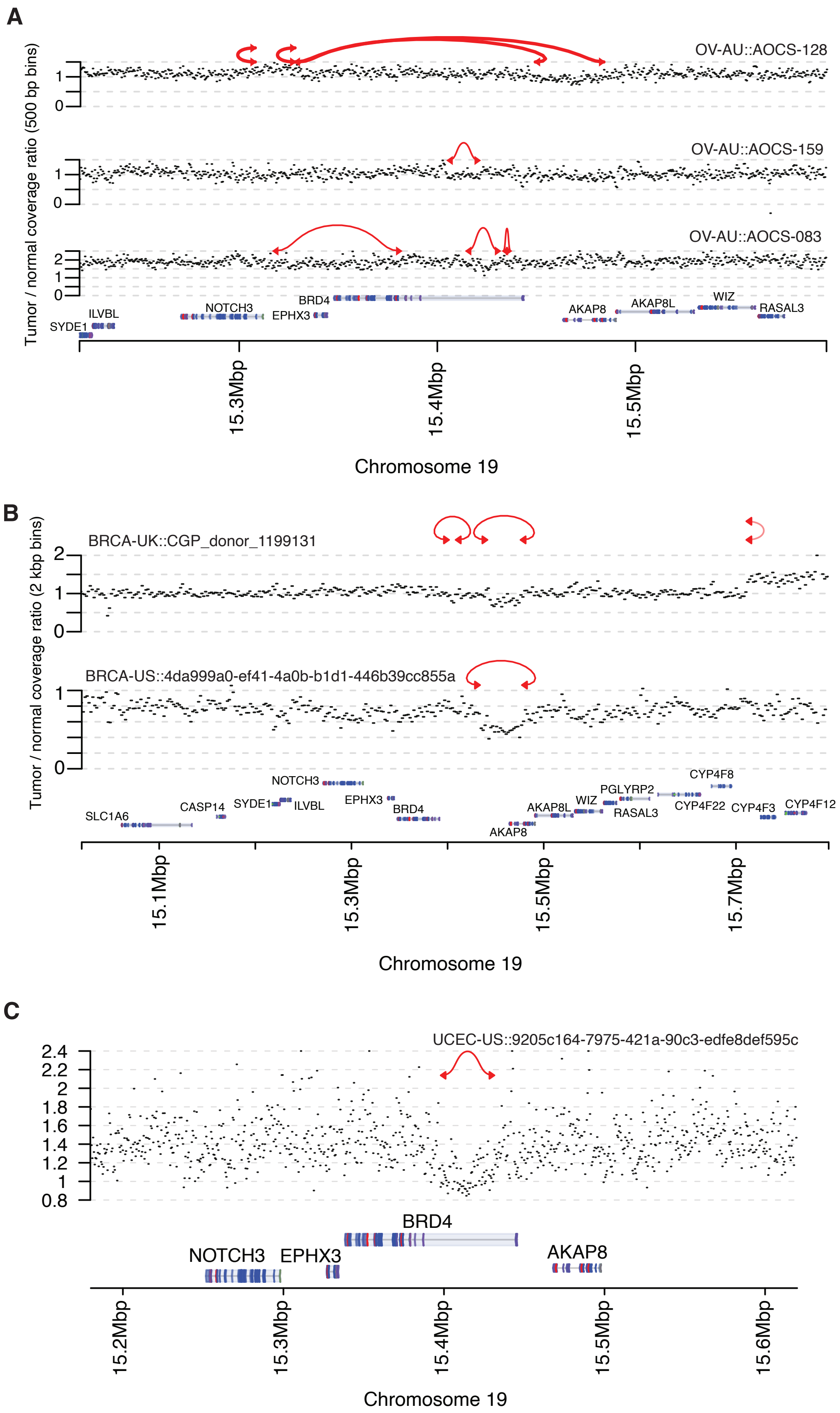
